# Proximity-dependent proteomics of the *Chlamydia trachomatis* inclusion membrane reveals functional interactions with endoplasmic reticulum exit sites

**DOI:** 10.1101/285106

**Authors:** Mary S. Dickinson, Lindsey N. Anderson, Bobbie-Jo M. Webb-Robertson, Joshua R. Hansen, Richard D. Smith, Aaron T. Wright, Kevin Hybiske

## Abstract

*Chlamydia trachomatis* is the most common bacterial sexually transmitted infection, responsible for millions of infections each year. Despite this high prevalence, the elucidation of the molecular mechanisms of *Chlamydia* pathogenesis has been difficult due to limitations in genetic tools and its intracellular developmental cycle. Within a host epithelial cell, chlamydiae replicate within a vacuole called the inclusion. Many *Chlamydia*–host interactions are thought to be mediated by the Inc family of type III secreted proteins that are anchored in the inclusion membrane, but their array of host targets are largely unknown. To investigate how the inclusion membrane proteome changes over the course of an infected cell, we have adapted the APEX system of proximity-dependent biotinylation. APEX is capable of specifically labeling proteins within a 20 nm radius in living cells. We transformed *C. trachomatis* to express the enzyme APEX fused to known inclusion membrane proteins, allowing biotinylation and pull-down of inclusion-associated proteins. Using quantitative mass spectrometry against APEX labeled samples, we identified over 400 proteins associated with the inclusion membrane at early, middle, and late stages of epithelial cell infection. This system was sensitive enough to detect inclusion interacting proteins early in the developmental cycle, at 8 hours post infection, a previously intractable time point. Mass spectrometry analysis revealed a novel, early association between *C. trachomatis* inclusions and endoplasmic reticulum exit sites (ERES), functional regions of the ER where COPII-coated vesicles originate. Pharmacological and genetic disruption of ERES function severely restricted early chlamydial growth and the development of infectious progeny. APEX is therefore a powerful in situ approach for identifying critical protein interactions on the membranes of pathogen-containing vacuoles. Furthermore, the data derived from proteomic mapping of *Chlamydia* inclusions has illuminated an important functional role for ERES in promoting chlamydial developmental growth.

## Introduction

*Chlamydia trachomatis* is an obligate intracellular bacterium that infects mucosal epithelial cells of the endocervix and conjunctiva. It infects millions of people every year and is the etiological agent of ocular trachoma [1,2]. Although *C. trachomatis* infections are effectively treated with antibiotics, the majority of infections are asymptomatic and go untreated [3]. The consequences of long term infection can be severe, especially in chronically infected women that are at risk for developing pelvic inflammatory disease, ectopic pregnancy, or infertility as a consequence of infection [4]. Chlamydiae undergo a biphasic developmental cycle, characterized by transitions between infectious elementary bodies (EB) and metabolically active reticulate bodies (RB) [5,6]. During infection of a host cell, an EB attaches to and internalizes into a vacuole called the inclusion. Within the inclusion, EB–RB conversion and replication proceed, ultimately followed by asynchronous conversion to EB and exit from the host cell. Chlamydial growth within host cells is critically dependent on protein rearrangements on the inclusion membrane early during infection, and on extracting nutrients from the host cell. The mechanisms responsible for these processes are not well understood. The streamlined genome of *Chlamydia* necessitates a dependency on the host cell for many nutrients, yet some of these molecules cannot freely permeate the inclusion membrane and import systems have not been identified [7]. In addition, while much is known about chlamydial manipulation of host signaling, very little is known about the molecular processes necessary for *Chlamydia* to obtain nutrients from the host [6,8].

The inclusion membrane represents the major interface through which chlamydiae manipulate host cell function. In accordance with this, different chlamydial species encode 50-70 type III secreted inclusion membrane (Inc) proteins that are predicted to localize to the inclusion membrane throughout the chlamydial developmental cycle [9–14]. The Inc family are a signature genetic feature of chlamydiae; however, their lack of sequence similarity with proteins from other bacteria has largely precluded bioinformatic prediction of molecular functions and potential host interaction targets. Two major proteomic studies in recent years have greatly advanced our knowledge of candidate host proteins associated with the inclusion membrane and with specific Inc proteins [15,16]. These studies also highlighted molecular interactions that occur between the inclusion membrane and the retromer complex [15,16].

A comprehensive understanding of inclusion membrane modifications, and the host proteins recruited to the inclusion, has not been realized. Even less is known regarding the temporal dynamics of these interactions over the 48-72 hour *C. trachomatis* developmental cycle, and the factors that are critical for inclusion biogenesis. Molecular analysis of early inclusions has been particularly elusive due to their small size. Previous proteomic efforts provided major new insight into the inclusion membrane proteome; however they were unable to characterize protein compositions of early inclusions or identify temporal protein associations. The development of techniques to enable study of the inclusion membrane proteome under native conditions, at multiple stages of infection, would provide new opportunities for investigating *Chlamydia*–host interactions. To this end, we adapted the APEX system of proximity-dependent biotinylation for use in *C. trachomatis*, as a flexible tool for exploring host-pathogen interactions. APEX is an ascorbate peroxidase that catalyzes a reaction between biotin-phenol and hydrogen peroxide, forming a phenoxyl radical that rapidly forms a covalent bond with a nearby amino acid [17,18]. The labeling radius of APEX is thought to be less than 20 nm, and the reaction is carried out in living cells for only one minute; this allows highly spatially and temporally resolved biotinylation of proteins in situ. In combination with mass spectrometry, APEX has been used to map the mitochondrial matrix, outer membrane, and inner membrane space, as well as the endoplasmic reticulum (ER) membrane and several other subcellular locations of mammalian cells [18,19]. Recently, APEX was used to study how protein interactions with G-protein-coupled receptors change after activation, highlighting its experimental utility for studying protein interaction dynamics [20]. We engineered a *C. trachomatis* strain that contained an Inc protein fused to APEX, and under the control of an inducible promoter. Using this strain, we delivered the APEX probe to the inclusion membranes inside cells infected with *C. trachomatis* at three stages of growth. Using quantitative mass spectrometry, we defined the proteomes of inclusion membrane interacting proteins at early, middle, and late stages of chlamydial development. Analysis of proteomic data for early inclusions showed a significant enrichment of endoplasmic reticulum proteins, in particular factors with established roles in the regulation of ER exit sites (ERES). We detail the recruitment of specific ERES factors to *C. trachomatis* inclusions, and we demonstrate that functional ERES are important for chlamydial developmental growth.

## Results

### Development of the APEX in situ proximity labeling system for *C. trachomatis*

We sought to develop the APEX system for identifying inclusion interacting proteins, with the primary goal of determining how these interactions evolve during *Chlamydia* infection. Previous mass spectrometry studies of the inclusion membrane were either done in the absence of infection, or only at a later stage of infection after mechanical manipulations [15,16]. These studies also used techniques that required lysing open the host cells and pulling down Incs or whole inclusions in an in vitro environment, which likely disrupted weaker or more transient protein-protein interactions. With the recent development of a transformation system for *Chlamydia*, a wide range of techniques are now possible [21,22]. Leveraging this advance, we infected cells with a *C. trachomatis* L2/434 strain engineered to express Inc–APEX fusion proteins, to enable the capture of inclusion membrane interacting proteins in live cells, at multiple times during infection. The APEX system is highly sensitive, has defined the proteomes of cellular compartments refractory to other techniques, and APEX enzymatic function was shown to remain intact when tagged to a chlamydial Inc protein [17,23,24].

We tested multiple Inc proteins (IncA, IncB, IncC, InaC, CT223) tagged to APEX and empirically found that IncB-APEX gave the most optimal combination of biotinylation levels and specificity on inclusion membranes (data not shown). We therefore chose to focus on IncB-APEX for the rest of our experiments. Although endogenously secreted IncB is known to localize at distinct microdomains in the inclusion membrane [14,25], overexpression of IncB caused it to distribute around the entire inclusion, thus allowing proteomic mapping of the broader inclusion membrane. Furthermore, IncB is constitutively expressed during infection, ensuring that any chaperones necessary for proper secretion will be present at any time points tested [26–28].

We transformed *C. trachomatis* L2/434 with a tetracycline inducible plasmid encoding IncB-APEX2 (Fig 1A). This genetic system allowed for tunable APEX expression and secretion at any time when the type III secretion system is active. The IncB-APEX2 fusion protein localized to inclusion membranes, with APEX exposed in the host cell cytosol. At specific time points, biotin-phenol and hydrogen peroxide were added to catalyze protein biotinylation, and resulting labeled proteins were pulled down using streptavidin and identified by mass spectrometry (Fig 1B). We confirmed that biotinylation required IncB-APEX2 expression, biotin phenol, and hydrogen peroxide (Fig 1C). Furthermore, the profile and depth of labeled proteins in cells infected with the IncB-APEX2 expressing *C. trachomatis* strain were distinct from cells infected with a strain expressing an untagged APEX2 that was not secreted (Fig 1C). Staining of infected, labeled cells with a fluorescent streptavidin probe demonstrated that biotinylated proteins were specific to the inclusion membrane at 8, 16 and 24 hours post infection (hpi) (Fig 1D). Biotinylated proteins at 8 hpi manifested as distinct foci on inclusion membranes, perhaps as a consequence of IncB-APEX2 accumulation at active sites of type III secretion [29].

**Fig 1.**
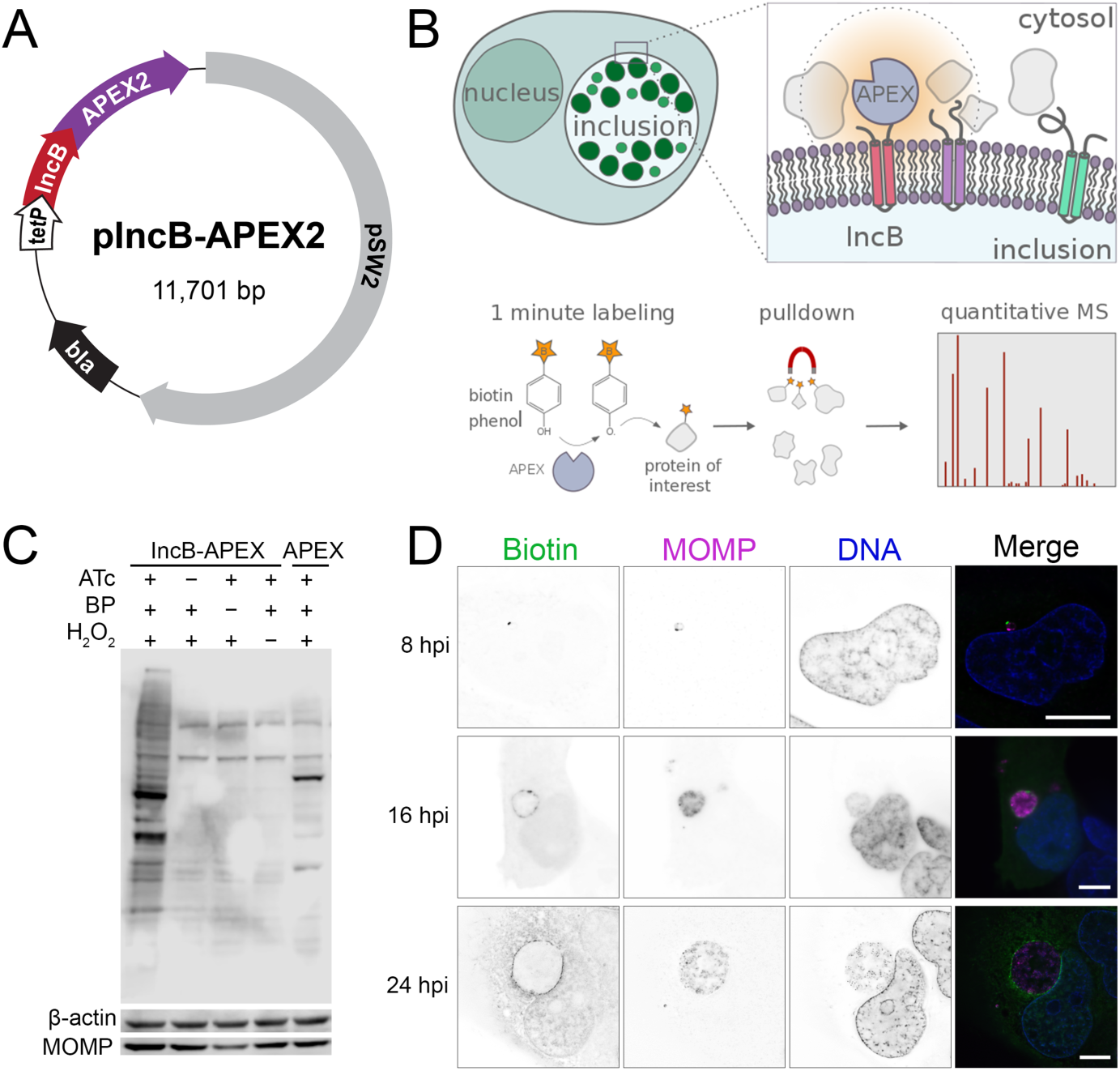
In situ proteomic labeling of the *C. trachomatis* inclusion membrane. (A) Plasmid used to transform *C. trachomatis* L2 and localize APEX to the inclusion membrane. IncB-APEX2 fusion expression was under the control of a tetracycline inducible promoter. (B) Schematic for APEX localization and biotinylation reaction. Cells were infected with *C. trachomatis* expressing IncB-APEX for 8, 16, or 24 hours, expression was induced by anhydrotetracycline, and incubation with biotin-phenol and hydrogen peroxide for 1 minute catalyzed biotinylation of proteins within 20 nm of APEX. Biotinylated proteins were enriched using streptavidin coated agarose resin and relative abundance estimated using mass spectrometry-based proteomics. (C) Western blot analysis of cells infected for 16 hours with *Chlamydia* expressing IncB-APEX (first four columns) or untagged APEX (last column). Blots were probed with streptavidin-HRP to detect biotinylated proteins. Anti beta-actin and anti MOMP antibodies were used as human and *Chlamydia* loading controls, respectively. BP, biotin-phenol; ATc, anhydrotetracycline. (D) Immunofluorescence microscopy of cells infected with *C. trachomatis* IncB-APEX and after 1 min inclusion membrane protein labeling at 8, 16, and 24 hpi. Representative images are shown. Biotin labeled proteins were identified by streptavidin-Alexa 488, *Chlamydia* were labeled with an anti-MOMP antibody, DNA labeled with DAPI. Single channel images are displayed in inverted grayscale. Merged panels display all three color channels. Scale bars = 16 μm.

### Identification of inclusion interacting proteins by quantitative mass spectrometry

HeLa cells were infected with IncB-APEX *C. trachomatis*, with 1 ng/ml anhydrotetracycline added at the start of infection to induce expression [30]. At 8, 16, or 24 hpi, infected cells were incubated with biotin-phenol for 30 min, and protein biotinylation was catalyzed by the addition of hydrogen peroxide for 1 minute. Following reaction quenching, cells were immediately pelleted and frozen for further processing. Control cells were treated identically to experimental cells, minus the addition of hydrogen peroxide. For each infection time point, biotinylated proteins were prepared from six biological replicates and six controls for enrichment and analysis by mass spectrometry.

Mass spectrometry of APEX-labeled samples identified 452 unique host proteins and 15 chlamydial proteins across the three time points analyzed. The presence of these proteins exhibited notable dynamics over the times tested, for example some only present at a single time point, and others maintained throughout infection (Fig 2A). Among host proteins recruited to *C. trachomatis* inclusions, 89 were significantly enriched at 8 hpi, 178 proteins at 16 hpi, and 396 proteins at 24 hpi. There were 37 proteins that maintained enrichment at all three time points (Fig 2A). Of the 15 chlamydial proteins labeled by IncB-APEX, 11 were annotated as Incs (Fig 2B, Incs highlighted in green). Although CT610 (CADD) does not have a canonical Inc structure, it has been shown to be secreted into the host cell by *Chlamydia* with an inclusion membrane distribution [31]. Of the remaining three *Chlamydia* proteins, two are found in high abundance in the bacteria and thus may be due to labeling of pre-secreted IncB-APEX. A complete list of proteomic data is contained in S1 Table.

**Fig 2.**
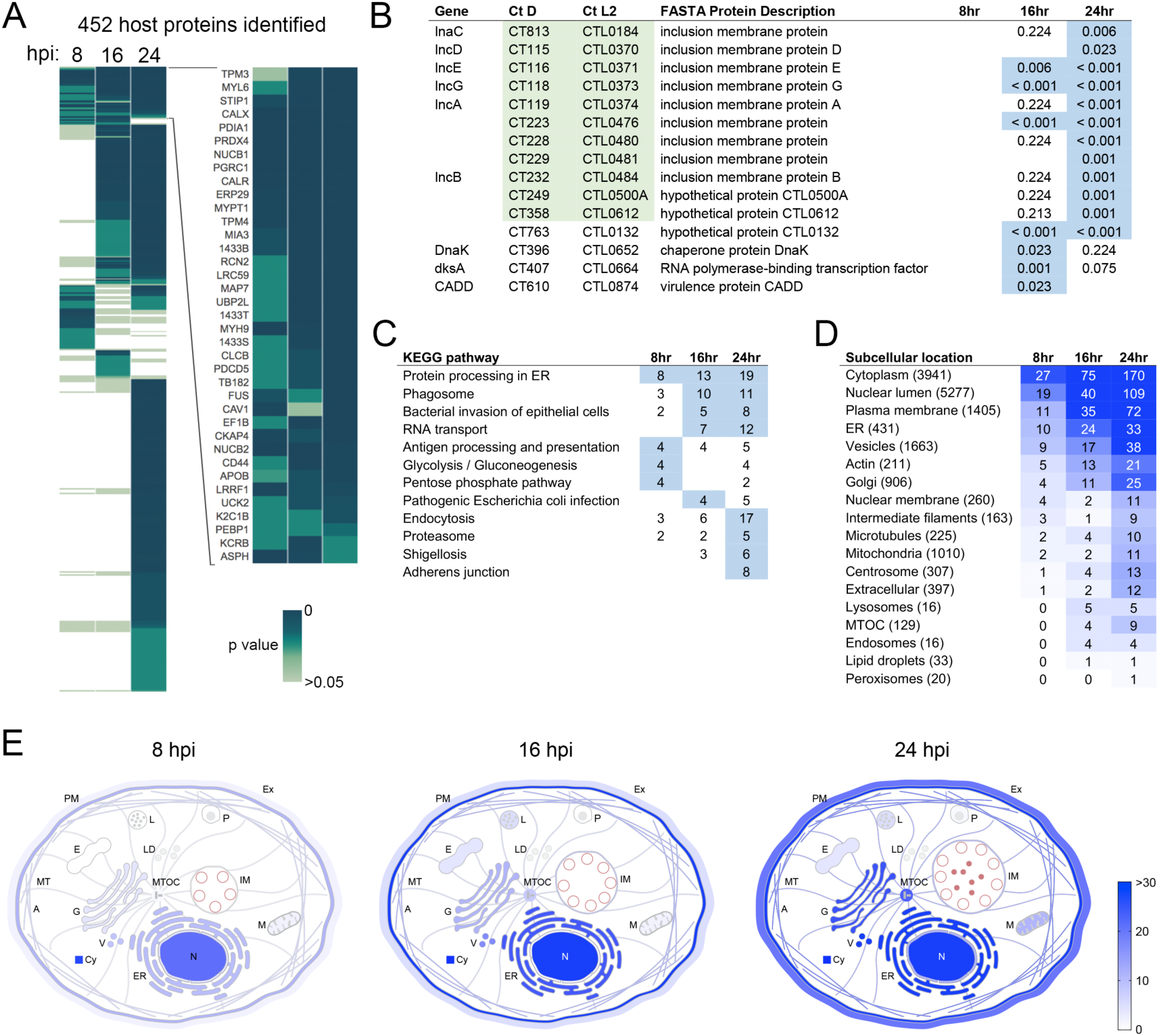
Global analysis of the inclusion membrane interaction proteome throughout the *C. trachomatis* developmental cycle. (A) Heatmap of inclusion membrane interacting proteome identified by APEX at 8, 16, and 24 hpi. Data from 6 replicate experiments per time point were averaged and compared against 6 replicate controls at similar times. Colors represent p-values, proteins not detected have no color. Enlarged section of heatmap shows proteins significantly enriched at all time points. Significance determined by T test or G test, p < 0.05. (B) *C. trachomatis* ORFs identified on the inclusion membrane, with p values displayed for their presence in 6 replicate samples for each time point. Locus tags highlighted in green represent annotated Inc proteins. Values highlighted in blue represent p values < 0.05. (C) KEGG pathway overrepresentation analysis of inclusion membrane proteome. Overrepresentation determined by hypergeometric algorithm with Benjamini Hochberg method for multiple test correction. Values highlighted in blue represent p values < 0.05. (D) Subcellular location enrichments of APEX identified proteins. Color intensity reflects the number of proteins for each location annotation pulled from the Human Protein Atlas database. Numbers in parentheses indicate the total number of reference proteins contained under that annotation in the Human Protein Atlas database. (E) Spatial distribution of inclusion interacting proteins from 8-24 hpi using Human Protein Atlas annotations and manual entry of Inc proteins. Color intensity indicates the number of proteins assigned to location annotations. A, actin filaments; Cy, cytosol; E, endosomes; ER, endoplasmic reticulum; Ex, extracellular/secreted; G, Golgi apparatus; IM, inclusion membrane; L, lysosomes; LD, lipid droplets; M, mitochondria; MT, microtubules; MTOC, microtubule organizing center; N, nucleus; P, peroxisomes; PM, plasma membrane; V, vesicles.

To identify novel *Chlamydia* interactions with the host cell, and to ensure that our approach labeled pathways and cellular components that *Chlamydia* is known to interact with, we analyzed proteomic data against annotation databases. Pathway overrepresentation analysis was performed using InnateDB with KEGG pathways, and the representation of subcellular locations in the data set was determined using the Human Protein Atlas database [32,33]. KEGG pathway analysis showed that many of the identified proteins were associated with cellular pathways known to play roles during *Chlamydia* infection, including ‘bacterial invasion of epithelial cells’ and endocytosis (Fig 2C). Proteins associated with ER processing were significantly enriched at all three time points. Proteins involved in glycolysis and the pentose phosphate pathway were significantly enriched at 8 hpi, suggesting an important role for recruitment and activation of these pathways by *C. trachomatis* early during infection. Analysis of subcellular localization annotations for IncB-APEX labeled proteins revealed a general spatial context consistent with the known perinuclear residence of the inclusion (Fig 2D,E) [8]. Ontology analysis indicated that early in *C. trachomatis* infection, at 8 hpi, inclusions acquired proteins normally localized to the cytosol, nucleus, plasma membrane, ER, and vesicles. These findings suggest an alignment of the chlamydial inclusion with the early secretory pathway, as opposed to usurping endosomal proteins. The latter category of proteins were not associated with early inclusions. Growth and maturation of the inclusion, at 16 and 24 hpi, was accompanied by a sustained enrichment of proteins associated with early inclusions, as well as an emergence of interactions with cytoskeleton associated proteins: actin, microtubules, centrosomes, the MTOC, and intermediate filaments. Inclusions are known to interact with multiple cytoskeletal components, and our data now provide evidence that these interactions are most heavily enriched with mature inclusions.

We used the STRING database to identify known molecular interactions between identified proteins, and used R to plot all proteins with at least one interacting partner (Fig 3A). This allowed analysis of the recruitment of multiprotein interaction complexes to the *C. trachomatis* inclusion. Highly interconnected protein communities were identified using the Louvain method of community detection. From this analysis, major protein networks emerged, encompassing innate immune signaling, the nuclear membrane, vesicular traffic, the cytoskeleton, nucleoside metabolism, and ER chaperones. Temporal analysis of these interactions revealed a dramatic recruitment of glycolytic proteins to early inclusions, for example aldolases, transketolase, pyruvate kinase, glucose-6-phosphate isomerase, and peroxiredoxin (Fig 3B). These data suggest a reliance of early inclusions on recruiting host glycolysis factors to the inclusion to support the early growth needs of the bacteria. Inclusion maturation was marked by a significant expansion of protein networks related to the MTOC and centrosome, innate immune signaling, proteasome regulation, and clathrin assembly (Fig 3B). Network analysis also allowed identification of host factors that may potentially interact with known host–Inc interactions, for example VAPA association with IncV [34] and the potential expanded interaction with S10AG and S10AE.

**Fig 3.**
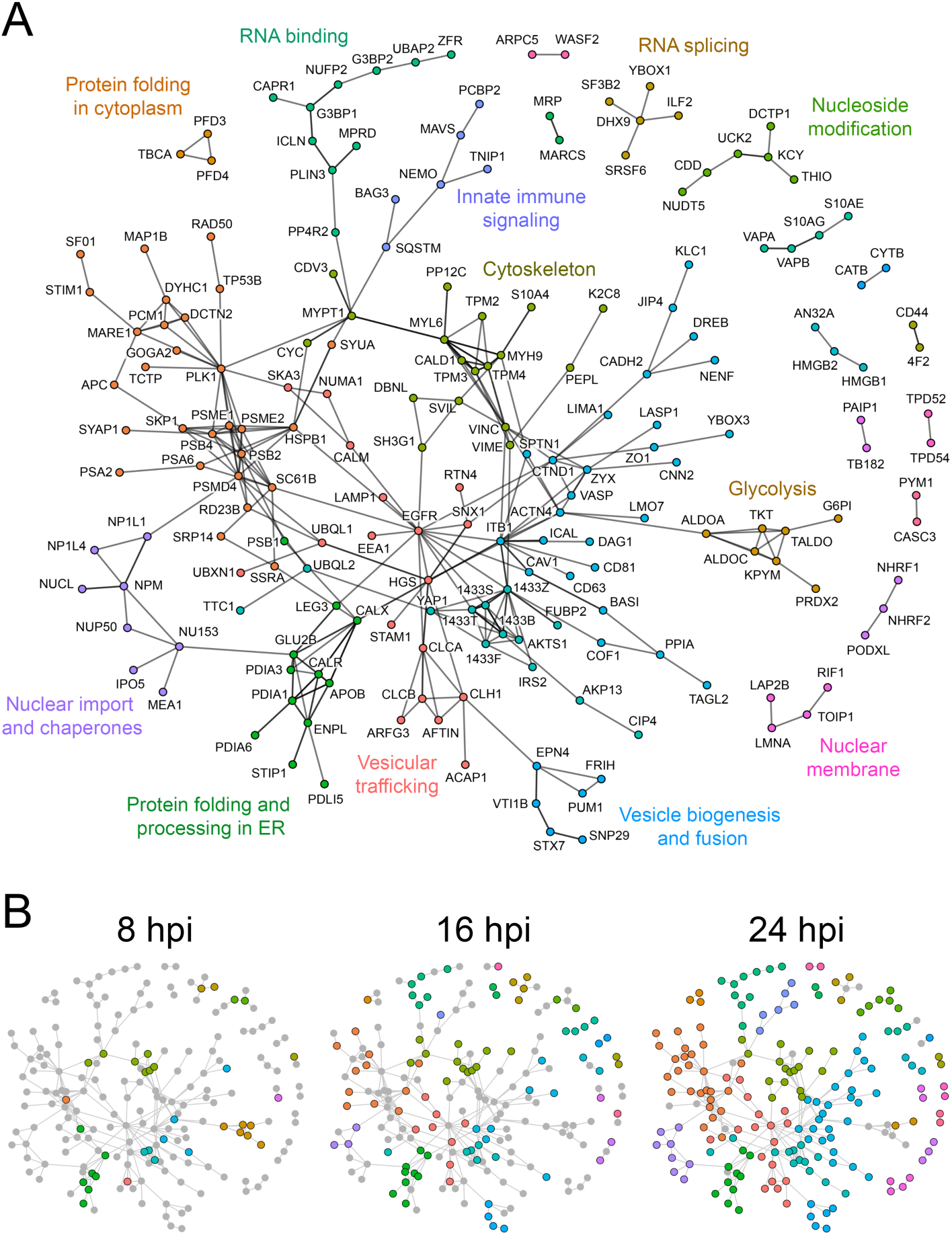
Network analysis of *C. trachomatis* inclusion membrane interacting proteins. (A) Interactions between host proteins identified by APEX proteomic labeling were obtained from StringDB [35], and visualization map was generated using R with the tidygraph package [36]. Colors represent more interconnected protein communities within the network. The interaction map in A represents the global extent of inclusion membrane interactions identified over the chlamydial developmental cycle. Edges between protein nodes represent interactions that were curated in StringDB either from published experiments or other databases. Proteins which do not have characterized interaction partners (i.e. single nodes) and translation related proteins were omitted for clarity. (B) Temporal dynamics of inclusion membrane protein interaction networks over the developmental cycle. Protein networks present in 8, 16, and 24 hpi proteomic data sets are represented by colored nodes. Gray nodes depict proteins present in the global interaction map (A) but absent from a specific stage of infection.

### Functional validation of targets recruited to early inclusions

We reasoned that proteins associated with early inclusion membranes might play important roles in mediating the biogenesis of the inclusion. To test this, and also validate the functional participation of inclusion interacting proteins towards *Chlamydia* developmental growth, we performed RNA interference on 64 of the 89 proteins present in the 8 hpi interaction dataset. HeLa cells were grown in a 96 well plate format, transfected with siRNA for 48 h, and infected with *C. trachomatis* L2 to determine the impact of protein depletion on bacteria growth. At 48 hpi, EB were harvested from siRNA treated cells and analyzed for inclusion forming units (IFU) on fresh cells. The results of the RNAi screen were that knockdown of 16 proteins (25%) resulted in a >1.5 fold change in infectious progeny formation, as compared to the mean IFU of the plate and a nontargeting siRNA control (Fig 4A). Because the screen used single oligonucleotide sequences per gene, these data likely underestimate the extent of 8 hpi interaction targets that functionally impact chlamydial growth. As a positive control, depletion of MAP1LC3B resulted in decreased *C. trachomatis* IFU, consistent with reported findings [37]. Overall, 10 protein knockdowns resulted in a >1.5 fold decrease in IFU (Fig 4A top), and 6 protein knockdowns led to a >1.5 fold increase in IFU (Fig 4A bottom). Proteins whose depletion resulted in decreased IFU tended to be functionally important at multiple stages of *Chlamydia* development (Fig 4A). In contrast, many of the proteins whose knockdown led to increased IFU production were more transiently recruited to inclusions, with enrichments primarily detected at early inclusions (Fig 4A). Collectively, the RNAi data show that early protein–protein interactions at the inclusion membrane functionally shape the inclusion in manner that impacts chlamydial growth.

**Fig 4.**
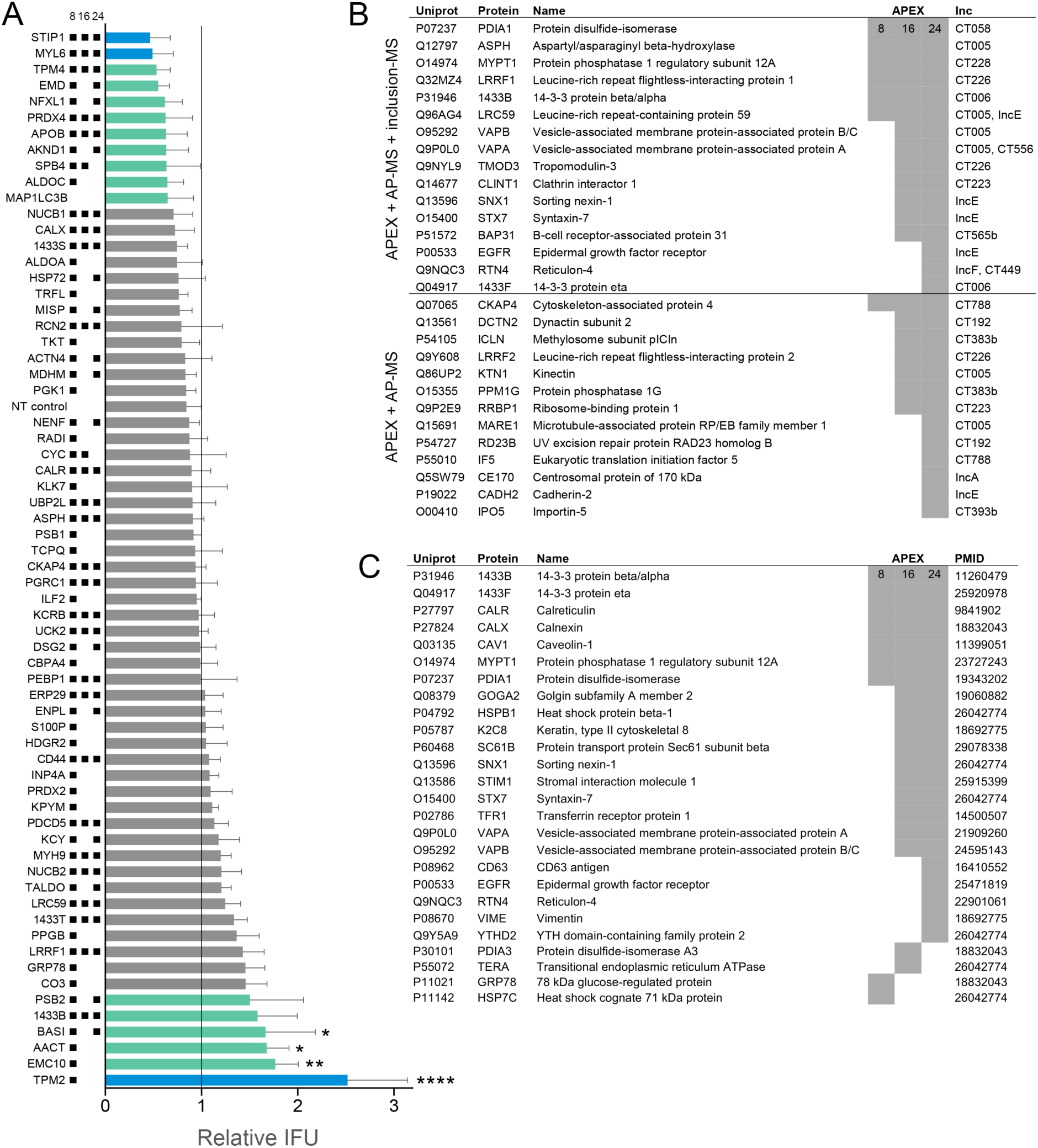
RNAi validation of inclusion interacting proteins and comparison to previous data sets. (A) IFU determination following RNAi depletion of 64 proteins identified from early (8 hpi) inclusions. Cells were transfected with siRNA corresponding to targets shown on the y-axis, infected with *C. trachomatis* L2, and harvested for IFU determination at 48 hpi. Bars, mean(SD); n = 3; IFU shown relative to mean IFU of plate. Black squares next to RNAi targets (y-axis) indicate which time points the protein was enriched in APEX mass spectrometry data. Teal bars correspond to at least 1.5 fold increase/decrease compared to the mean, blue bars are over 2 fold increase/decrease compared to the mean. Significance was determined by one-way ANOVA with Dunnett’s multiple comparisons test, comparing to mean infectivity of plate; *, p < 0.05; **, p < 0.01; ****, p < 0.0001. (B) Summary of all proteins shared between APEX data and two previous inclusion mass spectrometry data sets [15,16]. Lower table describes all proteins that overlap between APEX and AP-MS Inc-specific interactions [15]. Shaded boxes represent significant enrichment found by APEX. C. Summary of APEX identified proteins with reported microscopy-based associations with *Chlamydia* inclusions. PMID entries refer to the publications used to provide this evidence.

The data obtained for the dynamic in situ inclusion membrane interactome provided an opportunity to synthesize mass spectrometry data with those generated by previous studies, to develop a more comprehensive understanding of the protein networks recruited to the inclusion membrane. First, we compared our APEX data set to two previous mass spectrometry experiments that conducted analysis on harvested inclusions (labeled as ‘inclusion-MS’ in Fig 4B) [16], and affinity purified Incs expressed in 293 cells (labeled as ‘AP-MS’ in Fig 4B) [15]. Furthermore, proteins were manually cross-referenced against the literature to identify evidence-based reports of proteins recruited to inclusions or inclusion membrane proteins (labeled with PMID in Fig 4C). Overall, 16 proteins were identified by all three MS approaches to interact with inclusions, and our study now provides important temporal data for when these interactions occur during host cell infection (Fig 4B, upper half). Ten of the 16 proteins were previously reported to be recruited to inclusion membranes by immunofluorescence microscopy (Fig 4C). The remaining 6 proteins identified by all 3 MS approaches–ASPH, LRRF1, LRC59, TMOD3, CLINT1, and BAP31–are thus compelling candidates for further study in the context of *Chlamydia* infection. APEX labeling contained 60 additional proteins that were identified on 24 hpi inclusions by inclusion-MS (S1 Table) [16]. Thirteen proteins from APEX data correlated with Inc specific data obtained from AP-MS, thus adding in situ context to previously described molecular interactions (Fig 4B, lower half) [15]. Finally, APEX identified 26 proteins previously shown to be recruited to inclusions at a static stage of infection (Fig 4C). This substantiates the efficacy of the APEX approach and additionally provides important new temporal information for how these host targets are dynamically recruited to the inclusion membrane by *Chlamydia*.

### Recruitment of ERES proteins Sec16 and Sec31 to *C. trachomatis* inclusions

APEX proteomic analysis revealed a significant enrichment of ER associated proteins on inclusion membranes throughout infection. These findings are consistent with evidence for membrane contacts between inclusions and the ER [38,39]. APEX data showed that ER protein associations were present on early inclusions, at 8 hpi, and additionally highlighted an enrichment of markers for ER exit sites (ERES). ERES are specialized subdomains of the ER from which COPII coated vesicles bud and traffic to the ER-Golgi intermediate compartment (ERGIC) [40]. Sec16, TANGO1, TFG, and peflin, were identified on inclusion membranes by APEX, with TANGO1 present at all three time points (S1 Table). Sec16 is thought to act as a critical ERES scaffold, and directly stabilizes COPII subunits at ERES membrane regions [41]. TANGO1 recruits cargo to ERES through interactions with Sec16 and cTAGE5 [42,43], and TFG promotes COPII uncoating of vesicles prior to fusion with the ERGIC [44]. In support of an intimate association between ERES and inclusions, five members of the p24 family, proteins which regulate ERES organization and are packaged into COPII vesicles [45], were identified by inclusion-MS and AP-MS [15,16].

To validate these findings, we used immunofluorescence to investigate the spatial relationship between the ERES marker Sec16 and the inclusion membrane. In uninfected cells, Sec16 was distributed throughout the cytoplasm in small punctae, with a dense cluster of Sec16 in perinuclear ER membranes (Fig 5A). In *C. trachomatis* infected cells, Sec16 redistributed in a punctate pattern around the inclusion at 24 hpi (Fig 5A). Several Sec16 punctae adjoined inclusion membranes defined by IncA (Fig 5A, inset). Sec16 was also found to be recruited to inclusions at 14 hpi (S1 Fig). We further examined the distribution of Sec16 in live cells using HeLa cells transfected with Sec16-GFP and infected with a *C. trachomatis* strain expressing mCherry, and observed a similar recruitment of Sec16 to inclusions (S2 Fig). Next, we tested whether the COPII coat protein Sec31 similarly associated with inclusion membranes, as would be expected for functional ERES. Like Sec16, Sec31 was dramatically redistributed to inclusion membranes, with some Sec31 foci appearing to overlap with the inclusion membrane (Fig 5B, S1 Fig). Together, these data demonstrate that structural and regulatory ERES proteins are recruited to inclusion membranes. Previously, CERT was shown to be a key component of ER membrane contact sites with *C. trachomatis* inclusions [34,38]; CERT did not colocalize with ERES markers, however (S3 Fig), thus indicating that ERES are an additional type of ER interaction with chlamydial inclusions.

**Fig 5.**
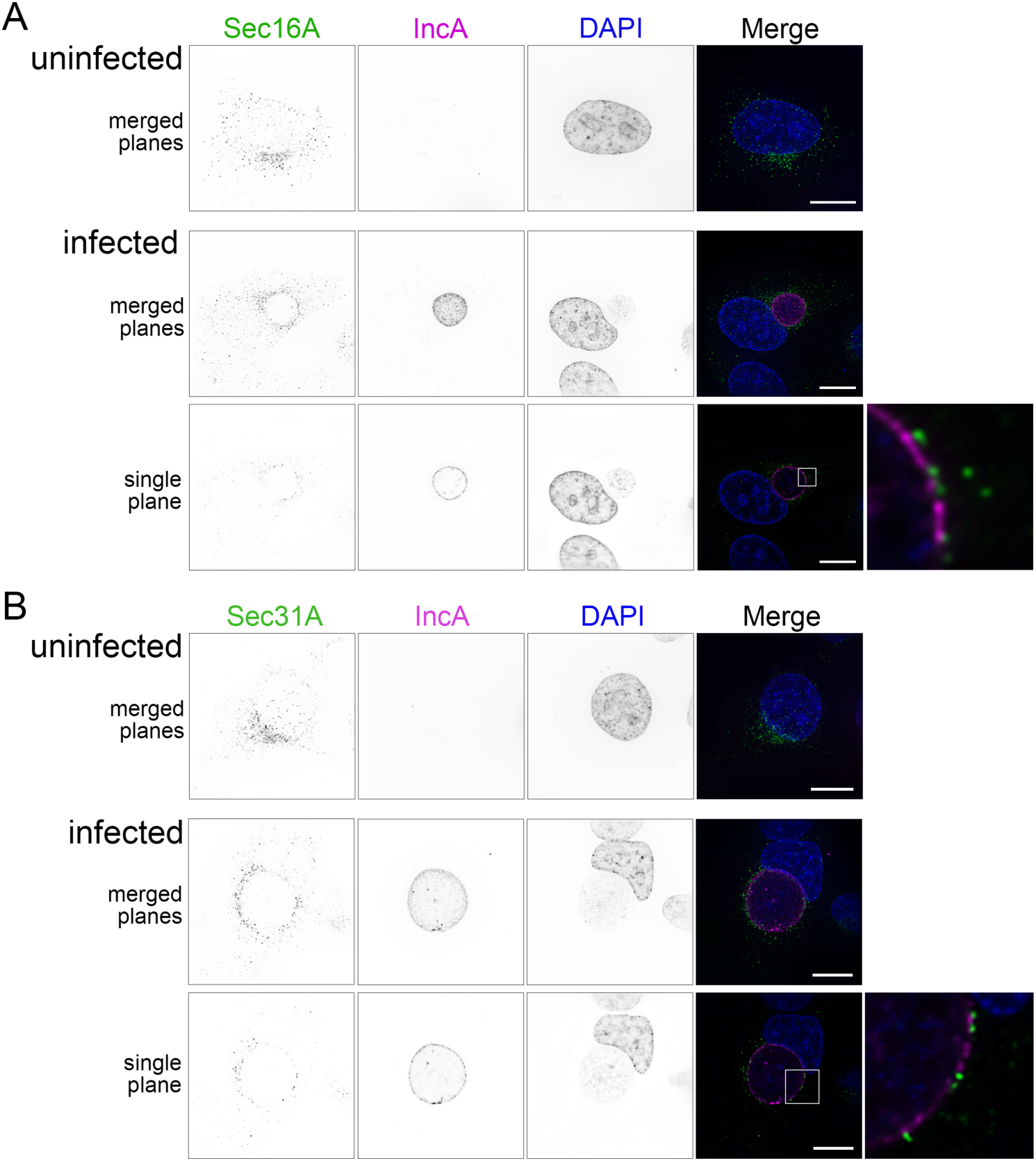
ERES proteins Sec16A and Sec31A are recruited to the *C. trachomatis* inclusion. (A) Immunofluorescence microscopy of cells showing cellular distribution of Sec16A (anti-Sec16A, first column, green in merge), *Chlamydia* IncA (anti-IncA, second column), magenta in merge), and DNA (DAPI, third column, blue in merge). (B) Distribution of the COPII outer coat protein Sec31A (anti-Sec31A, first column, green in merge), IncA, and nuclei. Single channel images are displayed in inverted grayscale. Merged panels display all three color channels. Protein distribution in uninfected HeLa cells are shown in the top row. Deconvolved images from cells infected with *C. trachomatis* L2 at 24 hpi are shown as a summation of z-series images (merged planes) or a single xy plane. Enlargements (far right) represent the regions marked with white boxes. Scale bars = 16 μm.

### Chemical inhibition of ERES cargo loading restricts *C. trachomatis* developmental growth

We next tested if functional ERES were required for *Chlamydia* infection, by using a specific inhibitor of ERES export, FLI-06 [46,47]. FLI-06 is a cell permeable, reversible inhibitor of ERES cargo loading and the early secretory pathway. First, we investigated how the localization of Sec16 and Sec31 were affected by inhibition of *C. trachomatis* infected cells with FLI-06. Treatment of infected cells for 4 h with FLI-06, from 20-24 hpi, abrogated the recruitment of both ERES proteins to inclusions, resulting in diffuse localization similar to that seen in uninfected cells treated with FLI-06 (Fig 6A,B). Similar effects were observed with live cells expressing Sec16-GFP (S2 Fig). These data indicate that Sec16 and Sec31 recruitment to inclusions are a functional consequence of COPII vesicle formation, and not a byproduct of their proximity to ER membranes.

**Fig 6.**
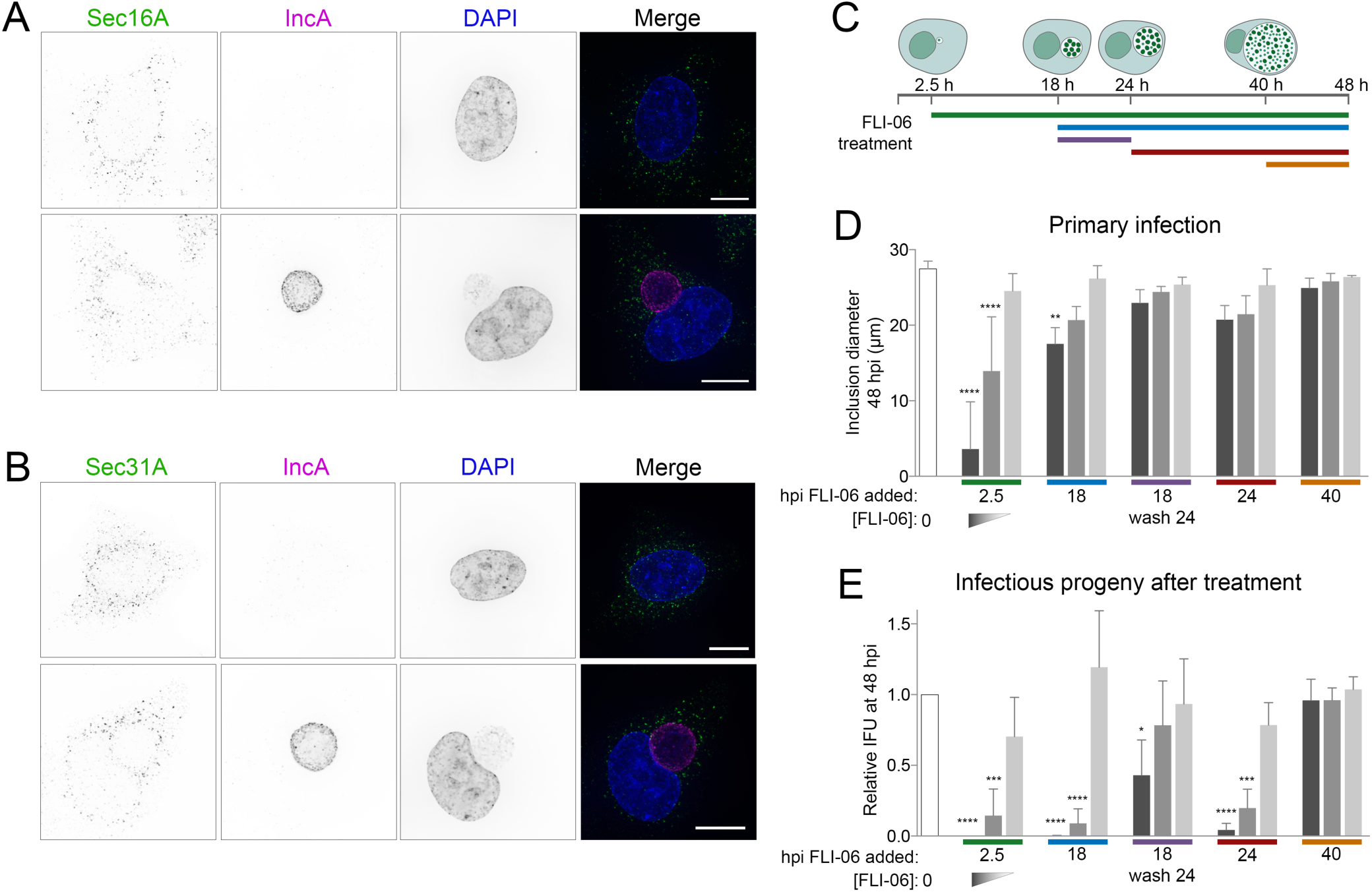
Inhibition of ERES cargo loading abrogates ERES recruitment to the inclusion and *C. trachomatis* developmental growth. Treatment of *C. trachomatis* infected cells with FLI-06 disrupted the recruitment of (A) ERES marker Sec16A and (B) COPII coat protein Sec31A to inclusion membranes. Immunofluorescence microscopy of cells showing cellular distribution of Sec16A or Sec31A (anti-Sec16A or anti-Sec31A, first columns, green in merges), inclusion membrane protein IncA (anti-IncA, second column), magenta in merges), and DNA (DAPI, third column, blue in merges). Representative deconvolved merged z-series images of uninfected cells are shown in upper panels, and cells infected with *C. trachomatis* L2 at 24 hpi are shown in lower panels. FLI-06 treatment for 4 h, from 20-24 hpi, resulted in Sec16A and Sec31A distribution similar to that of uninfected cells, and away from inclusion membranes. Scale bars = 16 μm. (C) Experimental design used to test the impact of ERES disruption on *Chlamydia* developmental growth. Colored bars mark the times when FLI-06 was applied to infected cells. All cells were harvested for IFU determination at 48 hpi. The effects of ERES inhibition were determined for (D) primary infection, through measuring the diameters of inclusions at 48 hpi, or (E) IFU production at 48 hpi. Bars denote the mean (n = 3; SD); white bars correspond to untreated (no FLI-06) controls; dark gray bars, 10 μM FLI-06; middle gray, 5 μM FLI-06; light gray, 1 μM FLI-06. Colored bars on x-axis correspond to the FLI-06 application key in C. Significance determined by one-way ANOVA with Dunnett’s multiple comparisons test, comparing to untreated control. *, p < 0.05; **, p < 0.01; ***, p < 0.001; ****, p < 0.0001.

To explore this further, we tested the effect of FLI-06 on chlamydial developmental growth. *C. trachomatis* infected cells were treated with three concentrations of FLI-06 at distinct stages of infection: 2.5, 18, 24, and 40 hpi. Following treatment, the effects of FLI-06 on primary infection and infectious progeny formation were determined by measuring inclusion diameter and IFU, respectively (Fig 6C-E). We measured a significant, dose-dependent reduction in inclusion diameter for infected cells treated starting at 2.5 or 18 hpi (Fig 6D, S4 Fig). Inhibitory effects were most pronounced with 10 μM of FLI-06, a concentration previously shown to block the recruitment of VSVG cargo to ERES [46,47]. The effects of FLI-06 were reversible, as treatment of infected cells at 18 hpi followed by washout at 24 hpi resulted in a recovery of inclusion growth at 48 hpi (Fig 6D). For cells treated with 10 μM FLI-06 at 2.5 hpi, fully formed inclusions at 48 hpi were rarely observed, indicating that early stages of *Chlamydia* and inclusion growth are reliant on COPII vesicle production from ERES. Two to 10-fold lower concentrations of FLI-06 resulted in dose-dependent phenotypes, thus strengthening the support that the effects of FLI-06 were biological.

The effects of FLI-06 on *C. trachomatis* IFU formation were more pronounced. Treatment of infected cells with 10 μM FLI-06 at 2.5 hpi resulted in no infectious progeny production, and a 99.8% or 95.6% reduction in *Chlamydia* IFU was observed in cells treated with 10 μM FLI-06 at 18 or 24 hpi, respectively (Fig 6E). Similar to the effects on primary infection, the impact of FLI-06 on *Chlamydia* growth was dose dependent. Cells pulsed with FLI-06 from 18-24 hpi partially recovered from the treatment, though IFU was reduced by 57.0% compared to untreated cells; this indicates that replication was reduced even during a 6 hour treatment. No effect on IFU production occurred when FLI-06 was applied to cells at 40 hpi, indicating that FLI-06 is not directly toxic to *Chlamydia* and that the effects of ERES on chlamydial inclusions were not on EB viability or infectivity. The potent effects of ERES disruption on inclusions at 18 hpi, when inclusions contain mostly RB, were consistent with an inhibitory effect that most heavily impacts RB growth. The application of FLI-06 at 18 hpi prevented the population of bacteria at that stage from converting into infectious EB by 48 h. Taken together, the data demonstrate that early *C. trachomatis* inclusions acquire vital cellular components from ERES-derived COPII vesicles for their complete developmental growth.

### Depletion of ERES regulatory proteins disrupts *Chlamydia* growth

We next sought to determine the specific ERES and COPII associated proteins responsible for providing factors critical for chlamydial growth. Using RNAi, we knocked down the expression of Sec16A and TANGO1, proteins which are required for efficient COPII transport [43]. We additionally knocked down cTAGE5 and Sec12; Sec12 is the guanine exchange factor for Sar1 GTPase, which in turn regulates COPII vesicle formation [48]. Sec12 also interacts with cTAGE5 at ERES, and it has been shown that cTAGE5 can recruit Sec12 to COPII budding sites [49]. We also depleted cells of Bet3, a key component of the TRAPP complex that functions in COPII vesicle tethering and fusion at the ERGIC [50].

HeLa cells were transfected with siRNA oligos and incubated for 48 hr prior to infecting with *C. trachomatis*. At 48 hpi, cells were lysed and chlamydial IFU was quantified on fresh HeLa monolayers. Knockdown of Sec12 and cTAGE5 resulted in a 45.1% and 39.6% reduction in IFU, respectively (Fig 7, S5 Fig). Surprisingly, Sec16A and TANGO1 depletion had no effect on IFU. Bet3 disruption also had no effect on IFU; however, this result was expected since TRAPP functions downstream of ERES and COPII vesicle formation, as a tethering complex on the ERGIC and cis-Golgi. Reasons for molecular specificity in the interaction between ERES and inclusion membranes are at this time unclear. The molecular mechanisms of ERES and ERGIC regulation are minimally defined, in particular the roles of cTAGE5 and TANGO1 at ERES. The apparent dispensability of TANGO1 and Sec16 for chlamydial growth is concordant with the finding that these proteins cooperate to recruit COPII components [43]. Complicating the molecular understanding of these processes are recent evidence for a cTAGE5/Tango chimeric protein that regulates cargo export from ERES, and poorly described roles for the cTAGE protein family in secretion [51,52]. FLI-06 blocks cargo recruitment to ERES, but unfortunately there are no known proteins involved in this step of COPII trafficking. All the genetic targets we tested were downstream of cargo recruitment, so it makes sense that the effect on IFU was not as dramatic. Thus, while it is clear that functional COPII vesicle production from ERES is critical for chlamydial growth throughout the developmental cycle, especially early during inclusion biogenesis, future efforts will need to resolve whether the basis of this interaction is to provide COPII vesicular cargo to inclusions, or non-vesicular nutrients from the ER, and to determine the identify of these ER-derived nutrients.

**Fig 7.**
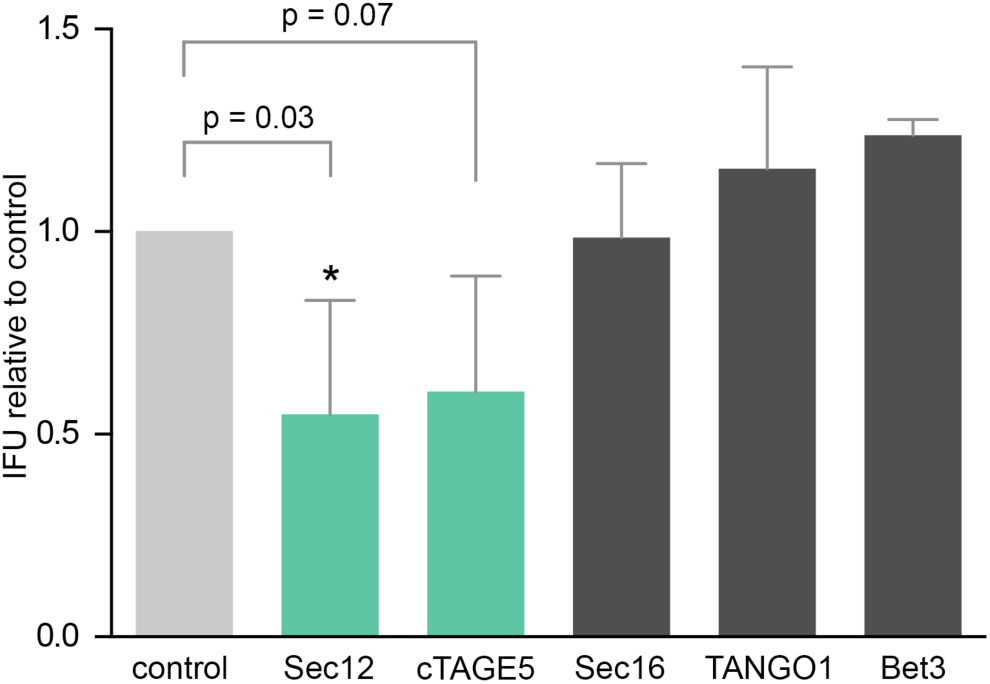
RNAi depletion of ERES regulatory proteins Sec12 and cTAGE5 disrupt *Chlamydia* growth. Cells were treated with siRNA oligonucleotides and incubated for 48 hours, then infected with *C. trachomatis* L2. At 48 hpi, cells were lysed and infectious *Chlamydia* EB from each sample group were tested for IFU by infecting new cells. IFU values were compared to scramble siRNA treated, infected cells. Significance determined by one-way ANOVA with Dunnett’s multiple comparisons test, compared to control IFU. Bars, mean (SD); *, p < 0.05; n = 3 (Sec16, TANGO1, Bet3), n=4 (Sec12, cTAGE5).

## Discussion

Establishment and maintenance of the inclusion is critical to *Chlamydia*’s ability to infect and grow within host cells. Bioinformatic analysis of the *C. trachomatis* genome predicts over 50 type III secreted Inc transmembrane proteins [7,9,11]; however, little is known about the broad spectrum of host proteins that are recruited to the inclusion membrane. Recent proteomic studies have provided a major, initial snapshot of proteins that comprise mature inclusions [16] and that associate with exogenously expressed Inc proteins [15]. However, we still lack an understanding of host factors recruited to inclusions in their endogenous host cell setting, how the inclusion membrane interactome changes throughout the developmental cycle, and in particular what signaling pathways are critical for inclusion biogenesis. To advance knowledge in these important unexplored areas, we developed the APEX proximity-dependent biotinylation platform for *Chlamydia*, to allow type III mediated expression of Inc proteins, fused to APEX, on the inclusion membrane. Using this approach, we obtained proteomic data for the dynamic recruitment of 452 host proteins to *C. trachomatis* inclusion membranes; this work strengthens existing proteomic datasets and additionally provides new insight into the *Chlamydia*–host interactions that shape the chlamydial intracellular inclusion niche. APEX proximity dependent proteomics represents a powerful tool for investigating host–pathogen interactions. In addition to IncB, we have tagged additional *Chlamydia* type III secreted effectors such as CT694, Tarp, IncA, IncC, InaC, and CT223. Previous work has demonstrated the efficacy of tagging IncF with APEX [24], and we urge the field to exploit this system to accelerate our understanding of the molecular functions of chlamydial type III secreted proteins.

A major finding of this study was the population of proteins assembled on early inclusions, as this subset is predicted to contain proteins important for regulating processes that shape inclusion biogenesis. Our data show that early inclusions, containing only a few bacteria, were enriched in proteins associated with the early secretory pathway and distinct cellular processes. In accordance with the need of chlamydiae to scavenge nutrients and energy from the host cell, proteins important for glycolysis were identified, including aldolases, transaldolase, transketolase, peroxiredoxins, and pyruvate kinase. In addition, a large number of ER proteins were proximity labeled by the inclusion membrane, for example protein disulfide isomerase, calreticulin, calnexin, endoplasmin, apolipoprotein B-100, STIP1, and EMC10. Finally, members of the 14-3-3 protein family, serpins, and several cytoskeletal proteins—alpha-actinin-4, emerin, myosins, MYPT1, tropomyosins—were found to interact with inclusion membranes. Depletion of many of these early association factors led to alterations in chlamydial growth, as measured by the production of IFU. We elected to perform a high throughput RNAi screen in order to test candidate proteins from the 8 hpi proteomic data. This approach contained single siRNA oligonucleotides for each target, and so it is likely that for some genes the absolute effects on IFU are underestimates. This is partly evident in the 35.2% decrease in IFU for MAP1LC3B knockdown cells, a ‘positive control’ target that we included based on a prior study which showed a 60% and 82% reduction in IFU relative to the control for two separate siRNA oligonucleotides [53]. Among proteins recruited to early inclusions, RNAi knockdown of tropomyosins yielded unexpectedly large and disparate effects on bacteria growth. Knockdown of TPM2 enhanced IFU by over 2 fold, while knockdown of TPM4 resulted in close to 2 fold reduced IFU. Although tropomyosins affect actin filament stability, the different forms are thought to be functionally distinct. Recruitment of tropomyosins to the inclusion could therefore be a strategy for stabilization of actin on the inclusion membrane, and this hypothesis is supported by the identified recruitment of TMOD3, tropomodulin-3, to midstage inclusions and Inc CT226. Furthermore, TPM2 expression was shown to be downregulated during infection [54], suggesting that *Chlamydia* may modulate TPM2 expression to benefit its growth. This result could explain why the knockdown of TPM2 allowed *C. trachomatis* to grow better, perhaps by decreasing TPM2 expression earlier in the developmental cycle.

Our study provides a third proteomic mapping of the inclusion membrane interactome, with each effort exploiting unique approaches and technological systems. We now have the opportunity to synthesize these proteomic datasets to derive a list of ‘high confidence’ interactions, and develop an understanding of multiprotein interactions that may occur with specific *C. trachomatis* Incs. Six proteins represented high confidence proteins recruited to early inclusions, three of which were previously shown to be recruited to inclusions: PDIA1 [55], ASPH, MYPT1 [56], LRRF1, 14-3-3β [57], and LRC59. By 16 hpi, seven additional high confidence proteins were found to interact with inclusion membranes across all three proteomic studies: VAPA [58], VAPB [59], TMOD3, SNX1 [16], STX7 [16], BAP31, and CLINT1. Finally, 24 hpi *C. trachomatis* inclusions were consistently enriched with three additional host proteins: EGFR [60], 14-3-3*ζ* [61], and RTN4 [39]. The high frequencies with which these proteins have been independently demonstrated to associate with chlamydial inclusions strongly suggests that their enrichments in proteomic data sets are not merely a byproduct of high protein abundance in cells. Interestingly, BAP31, ASPH, and LRC59 are normally associated with the ER; LRC59 additionally interacts with FGF, and *C. trachomatis* EB have been shown to interact with FGFR on the cell surface [62]. Furthermore, knockdown of early interacting high confidence proteins indicated that inclusion membrane associations for several—14-3-3β, LRC59, and LRRF1—have a functional impact on chlamydial growth. Depletion of these proteins resulted in increased IFU, suggesting that the targeting of these factors to inclusions is beneficial to the host cell. Another lens with which to interpret inclusion interactome data is towards resolving protein networks that are intimately associated with a particular Inc protein. For example, our data revealed the recruitment of DYHC1, PCM1, MARE1, MAP1B, and PLK1 on the inclusion membrane, and these interactions may be mediated through CT192, an Inc protein shown by AP-MS to also directly interact with DCTN2 [15]. Finally, our study indicated that chlamydial proteins constitute a small portion of the overall inclusion membrane proteome, as compared to host proteins. Only 11 Inc proteins were identified across all time points, suggesting that additional Incs are likely expressed at lower levels; Incs may also be difficult to detect compared to many more abundant host proteins. Another possibility is that most Incs are sequestered into heteromeric complexes, such that even with induced overexpression, IncB based proximity labeling was unable to access the full repertoire of Incs present on the inclusion membrane. Future efforts by the field should therefore focus on determining what extent of the inclusion membrane interactome is Inc-specific, and which Incs control the recruitments of host targets.

In this study, we also report the novel interaction between *C. trachomatis* inclusions and ER exit sites. The ERES associated proteins Sec16, TANGO1, peflin, and TFG, were identified by APEX as inclusion membrane interacting proteins, and follow up investigations confirmed the recruitment of Sec16 and the COPII coat protein Sec31 to the cytosolic surface of inclusion membranes. Inclusion–ERES interactions seem to be distinct from previously described IncD- and IncV-mediated ER–inclusion membrane contact sites (MCS) [34,38,39,59], since the ER membrane protein CERT, which interacts with IncD at MCS, did not colocalize with ERES proteins. In this regard, our findings strengthen the theme of inclusion membrane and ER interactions as playing important roles for inclusion biology and function during infection. Atlastin-3 and reticulon-4, which play roles in regulating ER morphology and structure [63,64], were also identified by our data and inclusion-MS [16]. In addition to ERES and COPII components, inclusion membrane interactions contain ERGIC-53 and VIP36 [16], two homologous proteins which are primarily localized to the ERGIC and function to regulate COPII vesicle fusion, and ER-ERGIC-Golgi syntaxins 5 and 18 [16]. We propose a model wherein *C. trachomatis* inclusions intimately interact with COPII mediated ERES to ERGIC vesicular traffic (Fig 8).

**Fig 8.**
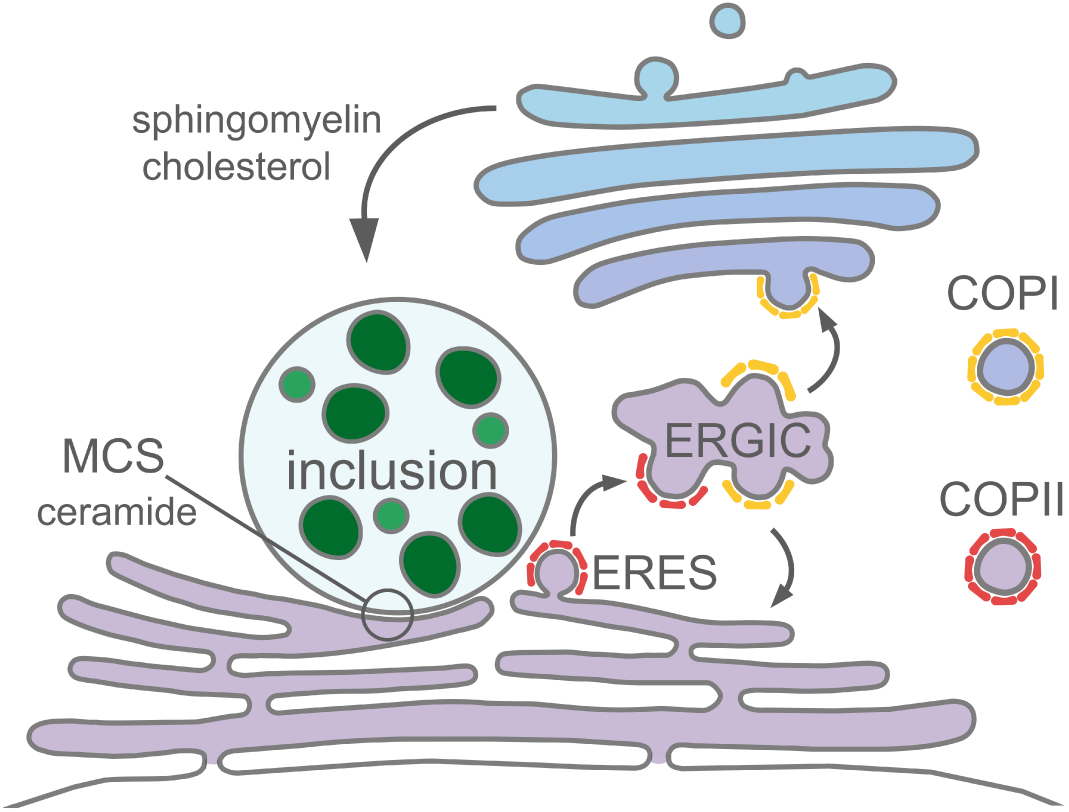
Model for interactions between the *Chlamydia* inclusion and the early secretory pathway. Our findings support a model wherein the inclusion membrane is situated in cells in close proximity to ERES domains. Normal COPII vesicle traffic from ERES are delivered to the ERGIC in mammalian cells, followed by COPI vesicle traffic to the Golgi complex. Inclusion–ERES interactions constitute a larger theme of physical and functional interactions between the inclusion and ER.

Importantly, functional disruption of ERES cargo loading using chemical and genetic approaches resulted in major defects on inclusion and chlamydial growth. Collectively, the functional and proteomic data show that complete developmental growth of *Chlamydia* requires efficient COPII vesicle production from ERES throughout the developmental cycle. Our findings for inclusion–ERES interactions, combined with the other nonredundant points of interaction between inclusion and ER, highlights the broad and critical need of *C. trachomatis* to extract lipids or other molecules from the ER. The ultimate unanswered question is then why *C. trachomatis* evolved to do this. At this time it is unclear what nutrients or metabolic benefit *Chlamydia* gains from interacting with ERES and potentially intercepting COPII vesicles. Data are inconclusive for indicating whether inclusion–ERES interactions represent COPII vesicles communicating with the inclusion membrane, or if these interactions consist of prebudding ERES complexes on the ER. One attractive nutritive benefit for *Chlamydia* is the acquisition of lipids, as supported by evidence that chlamydiae acquire phosphatidylcholine and other phospholipids from host cells [65,66], recruit lipid droplets to inclusions [67], redirect cholesterol to inclusions [68], and require functional host fatty acid synthesis machinery [67,69]. Disruption of many of these processes negatively impacted *Chlamydia* growth; however, the partial effects in every instance suggests that chlamydiae exploit numerous nonredundant mechanisms for acquisition of host lipids. *Chlamydia* is unable to synthesize phosphatidylcholine so it must be scavenged from the host cell [66], yet the mechanisms by which phosphatidylcholine is trafficked to inclusions are largely unknown [70]. Much of our understanding for ERES secretory mechanisms has come from studying the assembly of large cargo into COPII vesicles; for example procollagen, pre-chylomicrons, and very low-density lipoproteins [71]. For export of these bulky cargo, cTAGE5 and TANGO1 are essential. In our system, cTAGE5 disruption impaired IFU production, whereas TANGO1 knockdown had no discernible effect. An explanation for these findings is elusive, notably because only a preliminary functional characterization of these regulatory proteins exists. Interestingly, cTAGE5 and the TANGO-related gene MIA2 can form a fusion protein (TALI) which binds TANGO1 and facilitates the recruitment of apolipoprotein B containing lipid complexes to ERES [52]. Based on our proteomic data, apolipoprotein B is recruited to inclusion membranes throughout the developmental cycle. Finally, recent work has shown that starvation induces Sec12 and cTAGE5 relocation to the ERGIC followed by generation of autophagosome membrane precursors [72,73]. Future advances in unraveling ERES regulatory mechanisms in mammalian cells should fuel parallel efforts in determining how *Chlamydia* inclusions recruit and benefit from ERES protein interactions.

## Methods

### Antibodies and reagents

All reagents were purchased from Thermo Fisher Scientific (Rockford, IL) unless otherwise noted. Primary antibodies used in this study and their catalog numbers: Sec12 (PA5-53125), cTAGE5 (PA5-29515), Sec31A (Cell Signaling, Danvers, MA; 13466S), Bet3 (TRAPPC3; PA5-55841), Sec16A (Bethyl Laboratories, Montgomery, TX; A300-648A-M), TANGO1 (MIA3; Millipore Sigma, St. Louis, MO; SAB2700012), IncA (gift from Daniel Rockey), MOMP (Virostat, Westbrook, ME; 1621), CERT (PA5-28788). Secondary antibodies used: Donkey anti-Rabbit HRP, Donkey anti-Goat Alexa 594, Goat anti-Mouse DyLight 350, Goat anti-Mouse DyLight 594, Goat anti-Rabbit Alexa 488, Streptavidin Alexa 488.

### Cell culture and *Chlamydia* infections

HeLa 229 cells or McCoy cells (latter used for *Chlamydia* transformation) were grown in Roswell Park Memorial Institute 1640 (RPMI; Gibco) medium supplemented with 10% fetal bovine serum (HyClone) and 2 mM L-glutamine (HyClone) at 37°C, 5% CO2. *Chlamydia trachomatis* LGV L2 434/Bu was grown in HeLa 229 cells for 48 hours, then infected cells scraped into sucrose phosphate buffer (5 mM glutamine, 0.2 M sucrose. 0.2 M phosphate), lysed using bead bashing, and cell debris cleared by centrifugation at 300 × g for 10 minutes. Supernatant containing *Chlamydia* aliquoted and stored at -80°C. HeLa cells were infected with *Chlamydia* diluted in Hank’s buffered salt solution (HBSS; Gibco) at an MOI ∼1 for 2 hours at room temperature. Cells were washed with HBSS, then incubated at 37°C in RPMI.

### Plasmid constructs and generation of transformed *Chlamydia* strains

*Chlamydia* plasmid expressing IncB-APEX2 fusion was made in a modified version of the pASK-GFP parent vector provided by Scott Hefty [30], where GFP was replaced with IncB-APEX2 [18] and the mKate2 sequence was removed. Plasmid grown in dam-E. coli (C2925; NEB, Ipswich, MA) prior to *Chlamydia* transformation. Plasmid transformed strains of *C. trachomatis* L2 were generated using established procedures [21]. Three different dilutions (undiluted, 1:2, 1:10) of *C. trachomatis* L2 stocks were made in 50 μL calcium chloride buffer (20 mM Tris pH 7.4, 100 mM CaCl2). In the same buffer, 3 ug plasmid was diluted to 50 μL and added to diluted *Chlamydia*. The 100 μL *Chlamydia*–plasmid mixture was incubated for 30 minutes at room temperature. McCoy cells were trypsinized and diluted to a final concentration of 4 x 107 cells/ml in calcium chloride buffer. After 30 minute incubation, 100 μL diluted McCoy cells were added to *Chlamydia* and DNA mixture, and incubated for 20 minutes. In a 6 well plate, 100 μL McCoy cells, plasmid, and *Chlamydia* mixture were added to 2 mL medium. At 12-15 hours post infection, cells were washed and medium added containing 2.5 U/mL Penicillin G (Millipore Sigma) and 1 ug/mL cycloheximide. Cells were incubated for 48 hours, then *Chlamydia* passaged onto fresh McCoy cells. After first passage, cells were grown in medium containing 10 U/mL Penicillin G and 1 ug/mL cycloheximide. Transformants were passaged at least 2 more times, every 48 hours, until inclusions were apparent. Stocks of transformed *Chlamydia* were frozen at -80°C as described above.

### Biotin-phenol labeling in live cells

Biotin-phenol labeling of *Chlamydia* infected cells was adapted from described a protocol [74]. HeLa cells were grown in 8 well chamber slides (for immunofluorescence detection of biotinylation; Nunc Lab-Tek), or a T75 flask (for western blot and mass spectrometry), then infected with *C. trachomatis* L2 expressing IncB-APEX at an MOI of approximately 1. Cells were incubated in RPMI containing 1 ng/mL anhydrotetracycline (ATc; Acros Organics, New Jersey) and 1 ug/mL cycloheximide (Gold Biotechnology, St. Louis, MO). At 30 minutes prior to time point, 2.5 mM biotin-phenol (Iris Biotech, Marktredwitz, Germany) in RPMI was added to cells. At each time point, 30% hydrogen peroxide diluted to 100 mM working stock in Dulbecco’s phosphate-buffered saline (DPBS; Gibco) was added to cells at a final concentration of 1 mM, and allowed to incubate with gentle rocking for 1 minute at room temperature. Labelling solution was aspirated and cells were rinsed 3 times in quenching solution (10 mM sodium ascorbate, 5 mM Trolox, 10 mM sodium azide in DPBS). For immunofluorescence, normal fixation and staining protocols were followed. For western blot and mass spectrometry, cells were scraped into quenching solution and centrifuged for 5 minutes at 3,000 x g, 4°C. Cells were lysed by resuspending in ice cold RIPA buffer (50 mM Tris, 150 mM NaCl, 0.1% SDS, 0.5% sodium deoxycholate, 1% triton X-100, pH 7.5) with Halt protease inhibitor cocktail (Pierce) and quenchers (10 mM sodium ascorbate, 5 mM Trolox, 10 mM sodium azide). Lysate was incubated for 2 minutes on ice, then sonicated 3 x 1 second at 20% amplitude, and clarified by centrifugation for 10 minutes at 15,000 x g, 4°C. Part of the lysate was reserved for western blot, and remaining lysate for each sample was snap frozen until mass spectrometry processing.

### Protein purification for mass spectrometry

Protein concentrations for individual samples were measured by BCA. The samples were normalized to 2 mg/mL for enrichment. Labeled protein lysates were enriched with streptavidin agarose resin (Thermo Fisher Scientific, Rockford, IL). The resin was prepped for enrichment by placing the resin in a Bio-Rad chromatography column (Bio-Rad, Hercules, CA) on a vacuum manifold. The resin was washed with 0.5% SDS in PBS (1 mL, repeat 2×), 6 M urea in 25 mM ammonium bicarbonate (NH4HCO3) (1 mL, repeat 2×), and PBS (1 mL, repeat 4×). The resin was transferred to 4 mL cryovials using two 1 mL aliquots of PBS. An additional 0.5 mL of PBS was added to each tube followed by 1000 μg of protein (in 1.2% SDS in PBS). The total volume of each tube was set to 3.0 mL, giving a final SDS concentration of 0.2%. Tubes were rotated end over end for 4 hr at room temperature. Following streptavidin capture of biotinylated proteins, the solution was transferred into the Bio-Rad columns, and the solution was removed. The resin was washed with 0.5% SDS in PBS (1 mL, repeat 2×), 6 M urea in 25 mM NH4HCO3 (1 mL, repeat 2×), Milli-Q water (1 mL, repeat 2×), PBS (1 mL, repeat 8×), and 25 mM NH4HCO3 (1 mL, repeat 4×). The enriched resin was transferred to sealed 1.5 mL tubes using two 0.5 mL aliquots of 25 mM NH4HCO3. Samples were centrifuged at 10,500 x g, and the supernatant was discarded. 6M Urea was added to the resin for each sample followed by 100 mM TCEP (20uL) and place on a thermomixer for 30 min (1200 rpm at 37°C). After the samples were reduced, 200 mM iodoacetamide (20uL) was added to alkylate the proteins. The resin was placed back on the thermomixer for 45 min (1200 rpm at 50°C) and covered in foil. Following alkylation, the samples were returned to the Bio-Rad column and rinsed with PBS (1mL, repeat 8×) and 25 mM NH4HCO3 (1 mL, repeat 4×). The enriched resin was transferred to sealed 1.5 mL tubes using two 0.5 mL aliquots of 25 NH4HCO3. Samples were centrifuged at 10,500 x g, and the supernatant was discarded. Enriched biotinylated proteins were prepared for LC-MS/MS analysis. 25mM NH4HCO3 (200 μL) was added to the resin for each sample, along with trypsin solution. Resin solutions were placed on the thermomixer at 37 °C set at 1200 rpm set for overnight. Following trypsin digestion, the tryptic peptides were collected, and the resin washed once with 25 mM NH4HCO3 (150 μL). Volatiles were then removed from the combined tryptic peptide supernatant using a speed vacuum. The dried peptides were reconstituted in 25mM NH4HCO3 (40 μL) and heated for 10 min at 37 °C with mild agitation. To remove any solid particulates, samples were centrifuged at 53,000 x g for 20 min at 4 °C. From each ultracentrifuge vial was removed 25 μL for MS analysis. Samples were stored at -20 °C until analysis.

### Mass spectrometry and bioinformatic analysis of proteomic data

Biotinylated tryptic peptides were enriched and separated using in-house reverse-phase resin columns by LC and analyzed on a Thermo Fisher Velos Orbitrap MS as described previously [75]. Instrument data was acquired for 100 min, beginning 65 min after sample injection into the LC. Spectra was then collected from 400-2,000 m/z at a 100k resolution, following by data-dependent ion trap generation of MS/MS using the top six most abundant ions, a collision energy of 35%, and a dynamic exclusion time of 30 s for discriminating against previously analyzed ions. MS/MS spectra were searched using the MSGF+ algorithm and a tag-free quantitative accurate mass and time (AMT) tag approach for subsequent unique peptide to protein mapping using LC-MS peak feature detection, as described previously [76] with the following modifications. Identified features from each MS dataset were filtered on an FDR of less than or equal to 1%. Unique fragments, requiring a minimum of six amino acids in length, were filtered using an MS-GF threshold of ≤ 1 ×10-9, corresponding to an estimated false-discovery rate (FDR) <1% at a peptide level. Resulting relative peptide abundances, in replicate across 48 biological samples including control samples, were log transformed and the data was processed for quality control. Elimination of statistical outliers were confirmed using a standard Pearson correlation at a sample level [77]. Parameters for removing inadequate data for qualitative statistics required a minimum of two observations for a peptide across all groups to be compared quantitatively or identified in at least half the biological replicates for a given condition group using previously described methods [78]. Peptides were normalized using median centering, adjusting for overall differences in abundances across samples. Statistical comparisons were made between control and biotinylated groups at each time point and evaluated for quantitative differences using a standard 2-sample t-test and a qualitative difference (presence/absence markers) g-test. Additional evaluations were done in a similar manner for comparison of conditions across time.

For KEGG pathway analysis, UniProt identifiers of proteins were uploaded into InnateDB pathway analysis, then processed using the pathway overrepresentation analysis tool. The recommended settings were used for the analysis: hypergeometric algorithm, Benjamini Hochberg p value correction method. Only pathways with p < 0.05 after Benjamini Hochberg correction were listed.

To assign subcellular localization ontologies to proteins, the Human Protein Atlas database (version 18) was used. Subcellular localization data for all proteins was downloaded, and data for proteins in APEX data set was extracted. Only subcellular locations with enhanced, supported, or approved reliability scores were used. Proteins with more than one annotated location were counted for each location. Highly similar categories were condensed, for example nuclear speckles, nucleoli, nucleoli fibrillar center, and nucleoplasm were all counted as nuclear lumen.

Network analysis of APEX proteins was done using annotations from StringDB (version 10). Only interactions with evidence from experiments or databases were considered, with a confidence score of at least 0.700 (high confidence). Proteins with no interaction partners were excluded. Annotated interactions were downloaded for the proteins each time point, and this data was compiled into a list of all interactions at different time points using Microsoft Excel. To make the graph more readable, proteins involved in translation were excluded. This data was imported into RStudio software, and plotted using the tidygraph package [36]. Code details are provided in the supplemental methods. Graph layout was arranged using the Fruchterman Reingold algorithm, and subsets of closely interacting protein communities were detected using the Louvain method. General categories of some of the protein communities were assigned based on similarities between the UniProt descriptions of proteins in that particular community. If there were only 2 proteins in a group, or there was no consensus between the UniProt descriptions, no category was assigned.

To compare to the AP-MS [15] and inclusion-MS [16] protein lists, full data sets were obtained from the supplemental data of each study. Proteins were matched based on UniProt identifiers, and all three data sets were combined using RStudio and Microsoft excel (S1 table). For the AP-MS data, MIST scores are listed (closer to 1 is better). All Inc proteins that were found to interact with the host protein were listed, with the MIST score from the first listed Inc was retained. For APEX and inclusion-MS, p values are listed.

### siRNA and plasmid transfections

For siRNA transfections, HeLa cells were plated in 24 well or 96 well plates to 60-80% confluence. For 24 well plates, 50 μL Opti-MEM (Gibco) medium containing 5 pmol siRNA oligonucleotides and 1.5 μL Lipofectamine RNAiMAX was added to each well. Following transfection, cells were incubated for 48 hours at 37°C prior to infection or protein analysis by western blot. For 96 well plate experiments, knockdowns were done in duplicate, with 10 μL Opti-MEM containing 1 pmol siRNA oligonucleotides and 0.3 μL Lipofectamine RNAiMAX added per well. Oligonucleotides were purchased from Dharmacon, sequences and catalog numbers listed in supplemental methods. Dharmacon siGenome smartpools of 4 different oligonucleotide sequences were used for Sec16, Tango1, and Bet3 knockdowns. Dharmacon siGenome individual oligonucleotides were used for all other knockdowns.

For Sec16-GFP plasmid transfections, HeLa cells were plated in 4 well chamber slides (Nunc Lab-Tek), and infected with *C. trachomatis* L2. Immediately after infection, each well was transfected with 50 μL Opti-MEM containing 500 ng plasmid and 1.5 μL Lipofectamine 2000. Plasmid pmGFP-Sec16L was a gift from Benjamin Glick (Addgene plasmid # 15776) [79].

### Immunofluorescence and live cell microscopy

For immunofluorescence microscopy, cells were rinsed in HBSS, then fixed for 15 minutes in 3.7% paraformaldehyde in HBSS. Cells were rinsed 2 times in HBSS, then permeabilized with 0.5% triton X-100 for 15 minutes, and blocked in 1% bovine serum albumin (BSA) in PBS for 20 minutes. Samples were incubated with primary antibodies for 1 hour at room temperature in blocking buffer, then rinsed 3 times in blocking buffer, then incubated 45 minutes at room temperature with secondary antibodies in blocking buffer. DAPI was used to stain DNA, and added during the secondary antibody step. For biotin labeling, Streptavidin-Alexa 488 was incubated for 30 minutes at room temperature. For imaging of live cells, media was replaced with HBSS before imaging. Cells were imaged on a Nikon Ti-E inverted microscope and images were captured on a Hamamatsu camera controller C10600. Images were processed using Volocity software (PerkinElmer, Waltham, MA).

### Western blot analysis

For western blot analysis of biotinylated proteins [74], 6x protein loading buffer was added to lysates, samples were boiled for 5 minutes, followed by cooling on ice. 15 µl lysate was loaded and run on 10% SDS gel in Tris running buffer by SDS-PAGE. Proteins were transferred from gels onto Immobilon PVDF membrane (Millipore Sigma) with a Pierce G2 fast blotter in Pierce 1-Step transfer buffer, then blocked with 3% BSA in TBST overnight at 4°C. Blots were incubated with streptavidin-HRP in 3% BSA in TBST for 30 minutes at room temperature, washed 4 x 5 minutes in TBST, incubated with chemiluminescent substrate (Li-Cor 92695000, Lincoln, NE) for 5 minutes, and imaged using a C-DiGit blot scanner (Li-Cor). For western blot analysis of RNAi knockdowns, cell pellets were lysed in ice cold RIPA buffer (50 mM Tris, 150 mM NaCl, 0.1% SDS, 0.5% sodium deoxycholate, 1% triton X-100, pH 7.5) with Halt protease inhibitors (Pierce) for 30 minutes on ice, with vortexing every few minutes. Lysates were centrifuged for 20 minutes at 14,000 x g, and supernatant added to 4x laemmli buffer (Bio-Rad, Hercules, CA). Lysates were run by SDS-PAGE on 5-15% or 5-20% mini-PROTEAN TGX stain free gels (Bio-Rad), transferred to Immobilon PVDF membrane (Millipore Sigma), blocked for 1 hr in 5% milk-TBST, and labeled with antibody and digitally imaged as described above.

### Inclusion forming unit analysis

Infected HeLa cells were lysed at 48 hpi by incubating in water for 20 minutes followed by pipetting to disrupt cells. Serial dilutions of lysate were plated onto fresh HeLa monolayers in a 96 well plate. At 24 hpi, cells were fixed and stained with DAPI and an anti-MOMP antibody. Using immunofluorescence microscopy, 10-15 fields per well were taken at 20x magnification. Inclusions and nuclei in each field were counted using the Fiji distribution of ImageJ [80]. The macro used to count inclusions and nuclei in images is provided in supplemental methods. The overall percentage of infected cells was used to compare IFU between conditions, and the relative IFU calculated for each experiment by comparing to control. For FLI-06 experiments, HeLa cells in 24 well plate were infected with C. trachomatis L2 at an MOI around 1. FLI-06 was resuspended in DMSO to a concentration of 20 mM and frozen at -80°C until use. For treating cells, FLI-06 in DMSO was diluted 1:2000 in normal cell medium to make a final concentration of 10 μM FLI-06 and serial dilutions were used to make medium with 5 μM and 1 μM FLI-06. At 2.5, 18, 24, or 40 hours post infection, medium was aspirated from different wells in the plate and replaced with pre-warmed medium containing three different FLI-06 concentrations. Control well medium was aspirated and replace with fresh pre-warmed medium. For the 18 hour time point, two wells per concentration were prepared, one of which was aspirated and washed with HBSS at 24 hpi, then incubated in fresh medium. Since the stability of FLI-06 was unknown, medium in wells treated at 2.5 hpi was refreshed at 24 hpi, with identical concentrations of FLI-06. At 48 hpi, primary infection was imaged on microscope and 10-15 brightfield images were taken per well at 20x magnification and inclusion diameter was measured using Volocity software. After images were taken, cells were lysed for IFU assay as described above. IFU was adjusted based on number of inclusions per field in primary infection due to differences in cell growth with the inhibitor.

### Statistical analysis

Statistical analysis was performed using GraphPad Prism software. Comparisons between IFU or inclusion diameter were analyzed using one-way ANOVA with Dunnett’s test for multiple comparisons. Graphs indicate mean and standard deviation, statistical significance is indicated as follows: *, p < 0.05; **, p < 0.01; ***, p < 0.001; ****, p < 0.0001.

## Author Contributions

Conceived and designed the experiments: MSD, ATW, KH

Performed the experiments: MSD, LNA, JRH

Analyzed the data: MSD, BWR, LNA, ATW, KH

Contributed reagents/materials/analysis tools: ATW, RDS

Performed the statistical analysis of the proteomic data: BWR, LNA, ATW

Wrote the paper: MSD, ATW, KH

## Acknowledgements

We thank Alex Merz, Liang Ge, Kota Saito, and Daniel Rockey for discussions related to ERES regulation and protein functions, and for critical evaluation of the manuscript.

